# Genomic Prediction of Depression Risk and Resilience Under Stress

**DOI:** 10.1101/599456

**Authors:** Yu Fang, Laura Scott, Peter Song, Margit Burmeister, Srijan Sen

## Abstract

Advancing our ability to predict who is likely to develop depression in response to stress holds great potential in reducing the burden of the disorder. Large-scale genome-wide association studies (GWAS) of depression have, for the first time, provided a basis for meaningful depression polygenic risk score construction (MDD-PRS). The Intern Health Study utilizes the predictable and large increase in depression with physician training stress to identify predictors of depression. Applying the MDD-PRS derived from the PGC2/23andMe GWAS to 5,227 training physicians, we found that MDD-PRS predicted depression under training stress (beta=0.082, p=2.1×10^−12^) and that MDD-PRS was significantly more strongly associated with depression under stress than at baseline (MDD-PRS × stress interaction - beta=0.029, p=0.02). While known risk factors accounted for 85.6% of the association between MDD-PRS and depression at baseline, they only accounted for 55.4% of the association between MDD-PRS and depression under stress, suggesting that MDD-PRS can add unique predictive power to existing models of depression under stress. Further, we found that low MDD-PRS may have particular utility in identifying individuals with high resilience. Together, these findings suggest that polygenic risk score holds promise in furthering our ability to predict vulnerability and resilience under stress.

## Introduction

According to the World Health Organization, depression is the leading cause of disease-associated disability in the world ^1^. As current treatments for depression only result in remission in a minority of cases and new treatments have been slow to emerge, the burden of depression, including suicide, has continued to grow.

In populations at high risk, prevention of depression may be an effective strategy. The U.S. National Academy of Medicine has highlighted the need to develop, evaluate, and implement prevention interventions for depression and other mental, emotional, and behavioral disorders ^2 3^. However, our current ability to predict those most at risk for depression is limited.

Genetic variation accounts for 30-40% of the population variation in unipolar depression risk ^4^. In the past few years, genome-wide association studies have, for the first time, identified a substantial number of variants associated with depression ^5 6^. However, no individual variants of moderate to large effect have emerged, with evidence indicating that risk for depression is distributed widely across genome ^7^. Because the effect size of identified depression variants is modest, any individual polymorphism has limited utility for risk prediction. Polygenic risk scores (PRS) provide a mechanism for aggregating the cumulative impact of common polymorphisms by summing the number of risk variant alleles in each individual weighted by the impact of each allele on risk of disease. In other disease phenotypes, PRS has shown utility in predicting disease. For instance, the PRS for cardiovascular disease substantially improves risk prediction for disease beyond known risk factors ^8^.

Prospective cohort studies are critical to evaluating the predictive power of PRS ^9^. With life stress accounting for 30-40% of the population variation in unipolar depression risk ^10^ and approximately 80% of depressive episodes are preceded by a major stressor ^11^, a promising strategy is assess depression PRS (MDD-PRS) to predict the development of depression under stress. However, the unpredictable nature of stress makes prospective studies of depression difficult. The first year of professional physician training, medical internship, is an unusual situation where the onset of stress can be reliably predicted. The prevalence of major depression increases 5-6 fold during internship, with a series of psychological and demographic factors predicting the development of depression during internship stress ^12 13^. Here, we utilize internship to assess the predictive power of MDD-PRS for depression under stress.

## Results

5,227 medical interns of European ancestry from the Intern Health Study were included for analysis. This sample consisted of 50.3% women and a mean age of 27.6 years. We measured depressive symptoms using the PHQ-9 questionnaire before internship year started (baseline) and every three months during the stressful internship year. Included participants completed the baseline survey and at least one quarterly survey (average number of follow up visits was 3.46, SD = 0.90). The participants had a mean PHQ-9 score of 2.5 (SD = 2.9) at baseline and 5.6 (SD = 3.8) during internship. 3.4% of the subjects met PHQ depression criteria (PHQ-9 score >= 10) at baseline, with the percentage increasing to 33.2% during internship (Table 1).

**Table 1.** Sample Characteristics (N = 5,227)

|  |  |
| --- | --- |
| **Years age, mean (SD)** | 27.6 (2.7) |
| **Female Sex, N (%)** | 2627 (50.3%) |
| **Personal History of Depression, N (%)** | 2434 (46.6%) |
| **Neuroticism Score, mean (SD)** | 21.2 (8.8) |
| **Early Family Environment Score, mean (SD)** | 11.7 (8.8) |
| **Baseline Depression Symptom Score (PHQ-9), mean (SD)** | 2.5 (2.9) |
| **Baseline PHQ depression, N (%)** | 176 (3.4%) |

### Association of MDD-PRS with PHQ-9 depressive symptom score

We used the summary statistics derived from the most recent Major Depressive Disorder (MDD) GWAS, a meta-analysis of the Psychiatric Genomics Consortium (PGC) MDD phase 2 and 23andMe, Inc., a personal genetics company ^7^, to calculate the MDD polygenic risk score (MDD-PRS) in our sample, including all genotyped common SNPs in the MDD-PRS calculation. The standardized MDD-PRS in intern subjects had a near-normal distribution (Figure 1).

**Figure 1.**
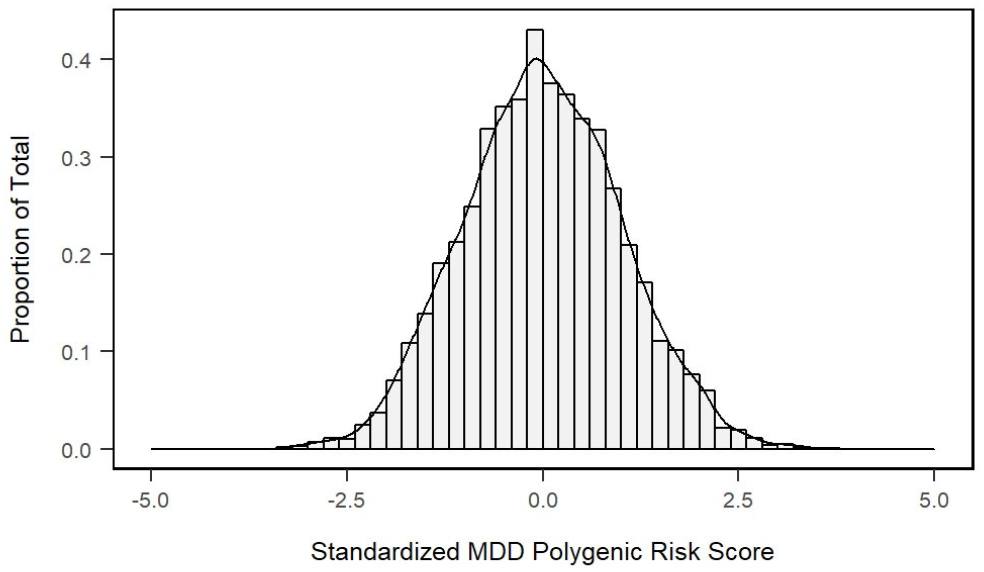
MDD-PRS Distribution. MDD-PRS has a near-normal distribution in Intern Health Study samples (n = 5,227). Represented on the x-axis, MDD-PRS was mean-centered and scaled to a standard deviation of 1.

To compare the predictive power of MDD-PRS on depression at baseline and during internship stress, we assessed the association between MDD-PRS and inverse-normalized PHQ-9 score under both conditions. After adjustment for age, sex and top 10 genotype-based PCs, one SD increase of MDD-PRS was associated with 0.052 higher PHQ-9 score at baseline (p = 1.7 × 10^−4^) and 0.082 higher PHQ-9 score during internship (p = 2.1 × 10^−12^) (Table 2, Figure 2a,b left plots). With both baseline and internship PHQ-9 scores included in the model, we found a significant interaction between MDD-PRS and internship stress status on PHQ-9 depressive symptom score (beta = 0.029, p = 0.023), indicating that the effect of MDD-PRS on PHQ-9 score was greater under internship stress than at baseline (Table 2).

**Table 2.** MDD Polygenic Risk Score Associations with PHQ-9 Depressive Symptom Scores

| Inclusion of known baseline risk factors* as covariates | Time Point of PHQ-9 score | Beta (SE) |  | p |  |
| --- | --- | --- | --- | --- | --- |
|  |  | MDD-PRS | MDD-PRS x internship stress Interaction** | MDD-PRS | MDD-PRS x internship stress Interaction** |
| Not included | baseline | 0.052 (0.014) | | $1.74 \times 10^{-4}$ | |
| | Internship | 0.082 (0.012) | | $2.11 \times 10^{-12}$ | |
| | baseline and internship | 0.051 (0.012) | 0.029 (0.013) | $3.38 \times 10^{-5}$ | 0.023 |
| Included | baseline | 0.008 (0.012) |  | 0.51 |  |
| | Internship | 0.040 (0.010) | | $7.52 \times 10^{-5}$ | |
|  | baseline and internship | 0.009 (0.011) | 0.029 (0.013) | 0.41 | 0.023 |

\* Neuroticism, Personal History of Depression, Early Family Environment
\*\* MDD-PRS x internship stress Interaction term is only available in the model including both baseline and internship data.

**Figure 2.**
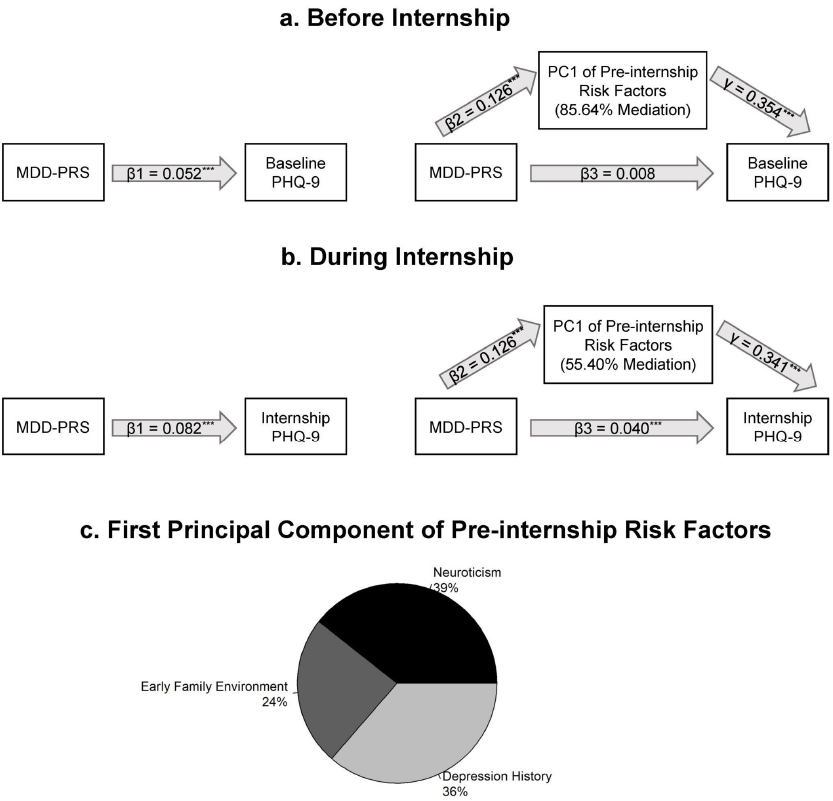
Associations of MDD-PRS and PHQ-9 Depressive Symptom Score and Mediations of the Associations by Known Risk Factors. **a)** Association before internship, without mediator (left diagram) and mediated by known baseline risk factors (right diagram). **b**) Association during internship, without mediator (left diagram) and mediated by known baseline risk factors (right diagram). **c**) Percentage of variance in first principal component explained by each risk factor (***: p<0.001)

To test the robustness of our findings to the set of variants used to calculate MDD-PRS, we also calculated MDD-PRS using LD pruned-imputed common SNPs. We observed slightly attenuated but significant associations between MDD-PRS and PHQ-9, both at baseline (beta = 0.042, p = 3.0 × 10^−3^) and during internship (beta = 0.070, p = 4.3 × 10^−9^) (Supplementary Table 1).

In addition to the quantitative PHQ score, we also utilized PHQ depression diagnosis as an outcome measure. In a logistic regression, we found no significant association between MDD-PRS and depression diagnosis at baseline (OR = 0.99, p = 0.88). In contrast, MDD-PRS was significantly associated with depression diagnosis during internship (OR = 1.17, p = 2.0 × 10^−7^). Parallel to the findings with PHQ-9 score, there was a significant interaction between MDD-PRS and internship status on PHQ depression diagnosis (OR = 1.29, p = 0.019) indicating the effect of MDD-PRS on depression prevalence was greater during internship than baseline.

Because the PGC/23andMe GWAS meta-analysis results were generated using European ancestry individuals, we restricted our main MDD-PRS analysis to the European ancestry subjects from our sample ^14^. To explore the predictive ability of European ancestry-based MDD-PRS in individuals of other ancestries, we assessed the association between MDD-PRS and PHQ-9 score in Intern Health Study participants of East Asian ancestry (n = 816) and South Asian ancestry (n = 595). We did not find and association between PGC/23andMe derived MDD-PRS and PHQ-9 scores in either the East Asian group (baseline beta = -0.011, p = 0.77; during internship beta = 0.012, p = 0.69) or the South Asian group (baseline beta = 0.028, p = 0.50; internship beta = 4.1 × 10^−4^, p = 0.99).

### Mediation of the association between MDD-PRS and PHQ-9 depressive symptom score by known risk factors

We conducted mediation analysis to quantify the proportion of the association mediated by three risk factors - neuroticism, personal history of depression and a stress early family environment - previously demonstrated to predict depression in both the general population, and training physicians, specifically ^12^. To capture the joint contributions of the three known depression risk, we performed principal component analysis and used the first principal component in our analysis (known risk factor-based PC). The contributions of neuroticism, personal history of depression, and early family environment in the first principal component were 39%, 36% and 24% respectively (Figure 2c). The known risk factor-based PC explained 85.64% of the association between MDD-PRS and PHQ-9 score at baseline but only 55.40% during internship (Table 3, Figure 2a,b right plots). The indirect effects of the known risk factor-based PC on PHQ-9 score were essentially the same at baseline (0.13 × 0.35 = 0.046) and during internship (0.13 × 0.34 = 0.044) (Figure 2a,b right plots), suggesting that the additional variance of PHQ-9 explained by MDD-PRS during internship was not mediated by the known risk factors but through other factors not included in the model.

**Table 3.**
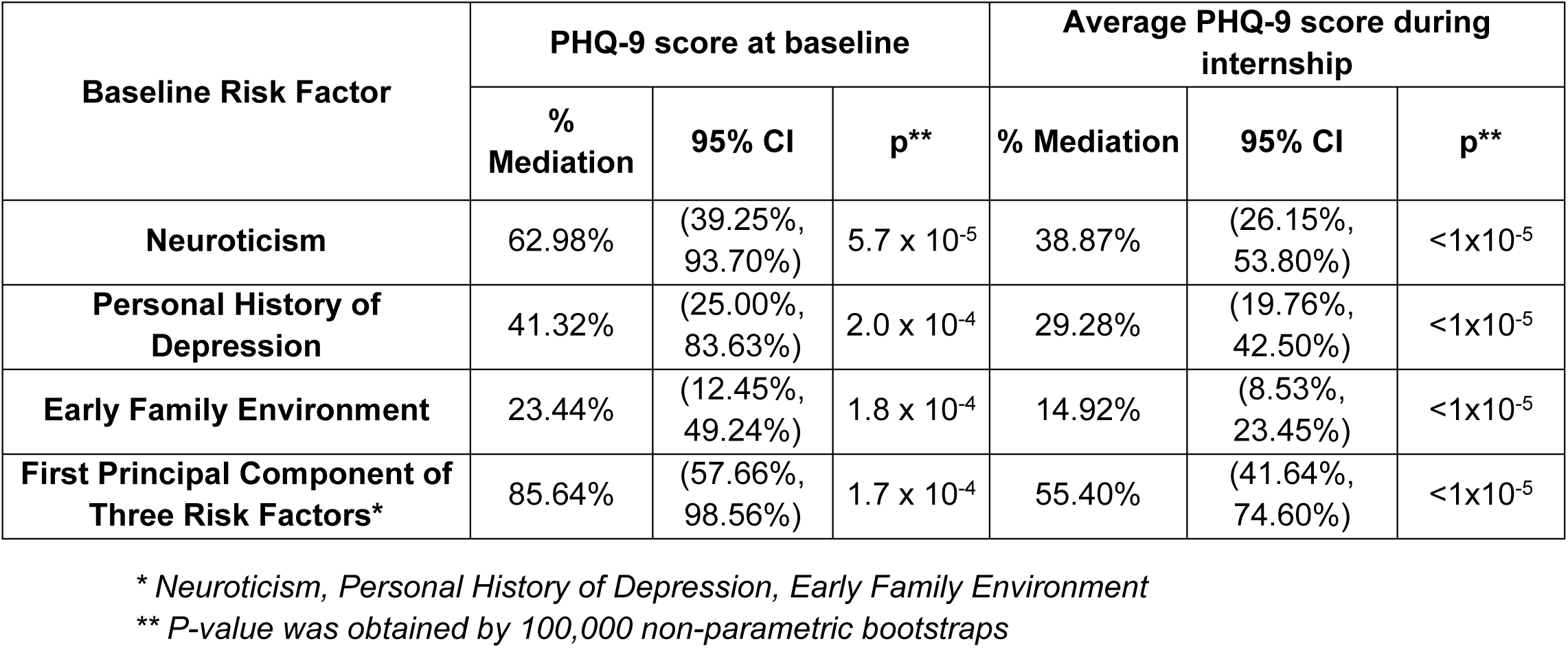
Mediation Test of Baseline Predictors on MDD Genetic Score Predicting PHQ-9 at Baseline and During Internship

### Differentiation of high risk/high resilience subjects

Khera and colleagues identified that individuals in the extreme high-tail of CVD-PRS distribution have several-fold higher risk of disease compared to other individuals ^8^. In order to assess if either extreme tail of the PRS distribution in our sample more effectively differentiated resilience versus susceptibility to depression during internship stress, we followed the approach of Khera and colleagues and divided our subjects into 40 quantiles from low to high MDD-PRS, each with 2.5% of the sample (n=131). Figure 3 displays baseline and internship PHQ depression proportion of each of the 40 MDD-PRS groups. The proportions of depressed subjects in the lowest MDD-PRS quantiles were below the trend line for association of MDD-PRS and depression, suggesting potential evidence for excess protective effects in the low tail of the MDD-PRS distribution. In contrast, subjects in the highest MDD-PRS quantiles were not above the trend line, providing little evidence of increased risk effect in the high tail of the distribution.

**Figure 3.**
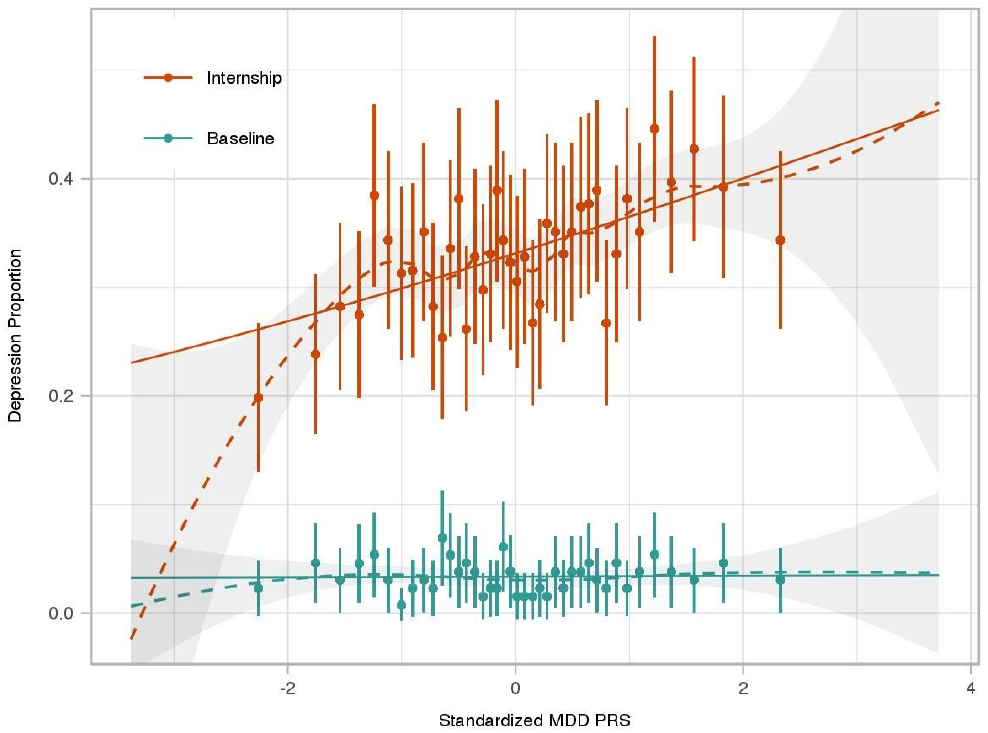
PHQ Depression Proportion by MDD-PRS Group. 5,227 Subjects from Intern Health Study were binned into 40 groups of 2.5% of subjects (n=131 per group) from low to high MDD-PRS (left to right). The 40 x-axis groups are defined by group-wise average standardized MDD-PRS. The proportion of subjects meeting criteria for PHQ depression at baseline (cyan dots) and during internship (orange dots) are plotted with 95% CI error bar. LOESS fitting line (dash line) shadowed by 95% CI and logistic regression fitting line (solid line) were applied to both baseline and internship plots. Optimal span parameter for LOESS regression was selected by generalized cross-validation method.

To quantitatively assess for a difference in depression risk prediction power between the lowest and highest tail, we serially dichotomized the sample using different MDD-PRS percentile cutpoints and compared the proportion of individuals with depression in the subjects above and below each of the cutpoints. For instance, subjects in the lowest 5% MDD-PRS distribution (low tail; n = 262) had lower rates of PHQ depression during internship (21.8%) compared to the remaining sample (33.8%) (OR = 0.54, 95%CI = 0.40 - 0.74, p = 3.7 × 10^−5^). In contrast, depression proportion of subjects with MDD-PRS scores in the top 5% (high tail; 36.8%) did not differ significantly from depression proportion of the remaining sample (33.0%; OR = 1.2, p = 0.18). Using a permutation test, we assessed if the odds ratio (OR) of the depressed subject for one tail was greater than the OR for the other tail (the reference sample for each OR being the subset of participants with lower MDD-PRS). For the 5% cutpoint, we found the low tail had a significantly larger OR than the high tail (p = 0.024, Table 4a). Similarly, using the PHQ-9 depressive symptom score, we found the test statistic for the low tail was significantly larger than that for the high tail (p = 0.038, Table 4b). These results indicate the lower 5% of the MDD-PRS distribution better differentiates depression risk and resilience compared to the upper 5% of the distribution.

When we tested more inclusive upper and lower tail cutpoints, we found that the differences in the proportion depression between the low MDD-PRS group and the remaining sample remained significant (for lower tail cutpoints at 10% (p = 1.6 × 10^−5^) and 25% (p = .003)) (Tables 4a). The differences in the depression proportion between the high MDD-PRS group and the remaining sample were also significant (for upper tail cutpoints at 10% (p = .003) and 25% (p = 3.7 ×10^−4^)) (Table 4a). The bottom OR and top OR were not significantly different for the 10% or the 25% cutpoint. We saw a similar pattern for PHQ-9 score (Table 4b), indicating no differences from what we would expect by chance for more inclusive cutpoints.

**Table 4.**
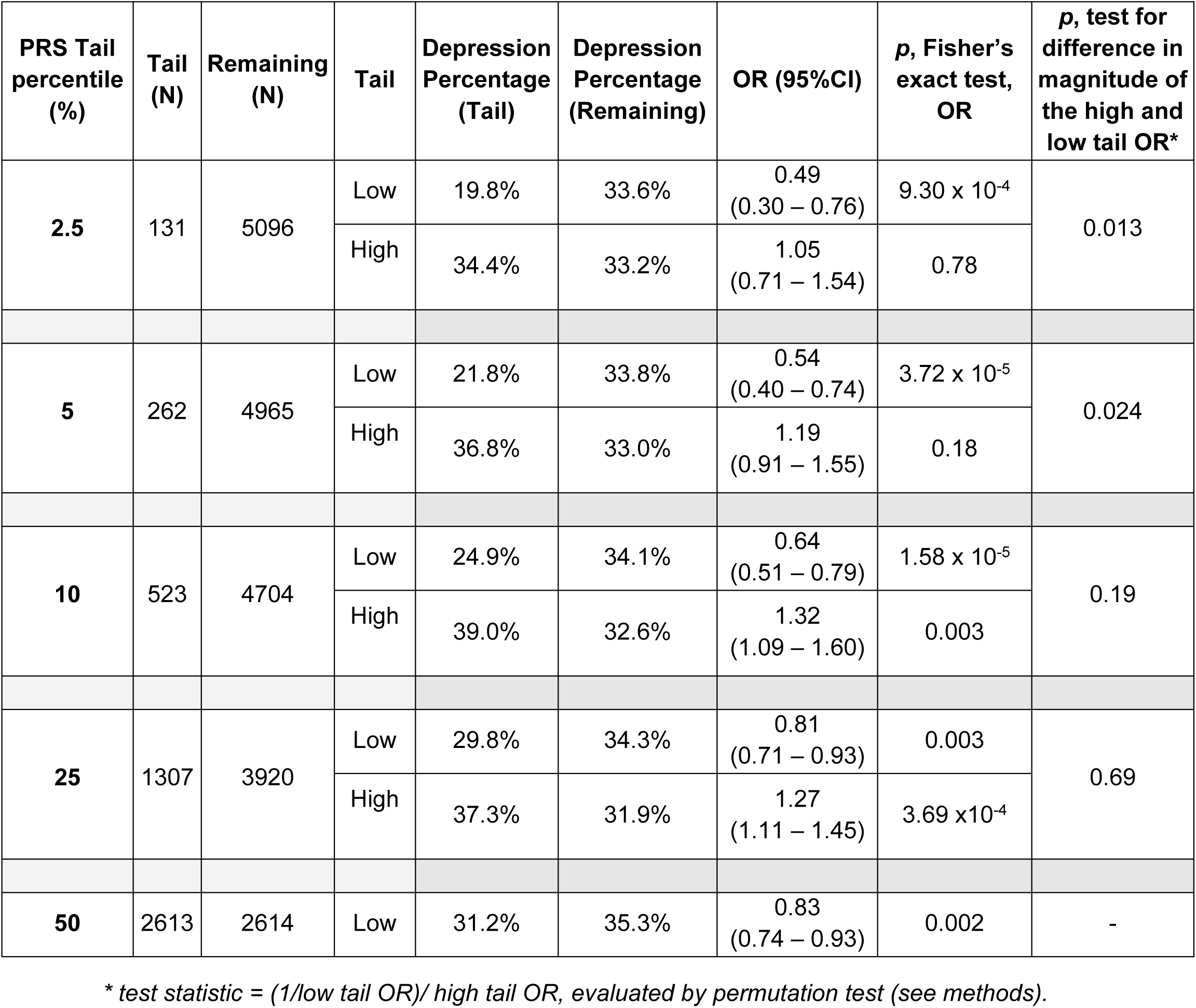
Comparison of Internship Depression in High and Low MDD-PRS Groups with the Remaining Sample across PRS Cutoffs Table 4a. Internship PHQ Depression in MDD-PRS Groups

**Table 4b.** Average Internship PHQ-9 Scores in MDD-PRS Groups

| PRS Tail percentile (%) | Tail (N) | Remaining (N) | Tail | PHQ-9 (Tail) | PHQ-9 (Remaining) | ΔPHQ, Tail - Remaining (95%CI) | *p*, two sample t-test, ΔPHQ | *p*, test of difference between high and low tail ΔPHQ* |
| --- | --- | --- | --- | --- | --- | --- | --- | --- |
| 2.5 | 131 | 5096 | Low | 4.40 | 5.61 | -1.21 (-1.79 ~ -0.62 ) | $7.42 \times 10^{-5}$ | 0.040 |
|  |  |  | High | 5.95 | 5.57 | 0.39 (-0.28 ~ 1.05) | 0.26 |  |
| 5 | 262 | 4965 | Low | 4.62 | 5.63 | -1.01 (-1.43 ~ -0.58) | $5.03 \times 10^{-6}$ | 0.038 |
|  |  |  | High | 6.01 | 5.55 | 0.46 (-0.06 ~ 0.97) | 0.080 |  |
| 10 | 523 | 4704 | Low | 4.83 | 5.66 | -0.83 (-1.16 ~ -0.51) | $7.18 \times 10^{-7}$ | 0.25 |
| | | | High | 6.16 | 5.51 | 0.65 (0.29 ~1.02) | $5.41 \times 10^{-4}$ | |
| 25 | 1307 | 3920 | Low | 5.16 | 5.72 | -0.55 (-0.78 ~ -0.32) | $2.45 \times 10^{-6}$ | 0.74 |
| | | | High | 5.99 | 5.44 | 0.55 (0.30 ~0.79) | $1.56 \times 10^{-5}$ | |
| 50 | 2613 | 2614 | Low | 5.35 | 5.80 | -0.45 (-0.66 ~ -0.24) | $1.92 \times 10^{-5}$ | - |

\* test statistic = ( - *t*-statistic<sub>low tail PHQ-9 - remaining PHQ-9</sub> ) - ( *t*-statistic<sub>high tail PHQ-9 - remaining PHQ-9</sub> ), evaluated by permutation test (see methods).

## Discussion

Building on the success of recent large-scale GWAS for MDD, this investigation utilizes a prospective cohort design to demonstrate that MDD-PRS is a significant predictor of future depression. Further, we find evidence that the association between MDD-PRS and depression is stronger in the presence of stress and that the additional predictive power of MDD-PRS under stress is largely independent of known risk factors for depression. Finally, we find that low MDD-PRS scores may have particular utility to identify individuals highly resilient to stress.

Our finding that MDD-PRS associates with depression during internship stress provides empiric evidence that the cumulative impact of common polymorphisms can produce meaningful risk prediction for depression. We find that individuals in lowest 2.5% of the PRS distribution in our sample have half the risk of developing depression compared to the rest of the sample. The predictive power of MDD-PRS will likely continue to improve with increasingly larger discovery GWAS studies. Based on the improving prediction profiles, individuals could be stratified into different strata of risk for transition to depression ^15^, with intensive prevention strategies targeted to high-risk individuals. For instance, in the population of training physicians, web-based CBT has been shown to be effective in the prevention of depression and suicidal ideation and could be targeted for prevention ^16^.

Under baseline, low stress conditions, the link between MDD-PRS and depression is largely explained by established risk factors measured in the study. In contrast, the established, measured factors only explained about half the association between MDD-PRS and depression under stress conditions. Understanding the outstanding mechanisms through which genomic predisposition leads to depression under stress could help to better elucidate how stress gets “under the skin” and exerts pathogenic effects. Further, effective risk predictors for depression will ultimately incorporate genetic variables with other established predictors. The finding that only about half the predictive power of MDD-PRS is mediated by established predictors of depression under internship stress suggests that genomics can add meaningful explanatory power to risk prediction from established factors.

We also find preliminary evidence that the overall association between MDD-PRS and depression under stress is driven disproportionately by the lower end of the PRS distribution. Specifically, while individuals with very low PRS scores are substantially less likely to become depressed relative to the rest of the sample, individuals with very high PRS scores do not show an analogously higher relative risk of depression. As a result, MDD-PRS may be better at identifying resilient individuals than at identifying those that are most at risk for depression under stress. With the relatively small number of subjects (N=131) in each of the PRS subgroups, these findings should be assessed in other prospective stress samples before drawing definitive conclusions. Resilience is a dynamic and active neurophysiological and psychological response to stress that is not merely the absence of vulnerability ^17^. Delineating the genomic factors that are protective against disease have informed genomically anchored medicines that assist in maintaining health ^18^. Identifying specific genes within the broad PRS that are driving resilience has the potential to drive new therapies for depression.

There are a number of limitations to this study. First, this study focused on physician training as a specific stressor. While the established predictors of depression are similar to predictors of depression in the general population, the predictive power of MDD-PRS should be explored in other prospective stress samples. Second, as the MDD-PRS incorporates input from across the genome, drawing definitive conclusions about underlying mechanism is not possible. Third, the polygenic scores described here were derived and tested in individuals of European ancestry only. Building up large enough samples to establish meaningful PRS for other ancestries is imperative to ensure that any benefits that follow from genomic medicine are not restricted to European ancestry populations. Lastly, the potential utility of genetic risk disclosure should be balanced against potential harm. Specifically, as PRS improves, protections should be put in place to ensure that prediction cannot be used to discriminate against at risk individuals. Reassuringly, our findings suggest that the MDD-PRS is better at identifying resilient individuals than individuals at most risk.

In summary, we find that MDD-PRS is a meaningful predictor of depression risk under stress. Future work should extend this work in multiple directions. First, the predictive power of MDD-PRS should be assessed in other prospective stress models, such as military stress and pregnancy. The similarities and differences in the genomic risk profiles across different types of stress would inform the extent to which future risk prediction work could be generalized across stressors or should be restricted to specific subtypes. Second, the finding that the percentage of the relationship between MDD-PRS and depression explained by established depression predictors decreased from 86% at baseline to 55% under internship stress suggests that there are important behavioral and psychological pathways from genomic risk to depression that are not fully explicated. Further elucidating those pathways could enhance our understanding of the pathologic effects of stress. Similarly, the finding that individuals with extremely low MDD-PRS scores are particularly resilient to depression under stress suggests that prospective study of the behavioral and biological responses to stress among individuals with low MDD-PRS scores could identify strategies to prevent depression and inform the incorporation of precision medicine into psychiatry.

## Methods

### Participants

The Intern Health Study is a multi-institutional prospective cohort study that follows training physicians through the first year of residency training (internship). Interns entering residency programs across specialties in the academic years from 2007 to 2017 were sent an email 2-3 months prior to commencing internship and invited to participate in the study. Subjects consented to participate in the study were given a $25 gift certificate after completing the baseline survey and another $25 gift certificate after completing the follow-up survey ^12 16^. The Institutional Review Board at the University of Michigan and the participating hospitals approved the study.

### Data Collection

The primary outcome of the study was depressive symptoms measured through the PHQ-9 (the 9-item Patient Health Questionnaire), a self-report component of the Primary Care Evaluation of Mental Disorders inventory. For each of the 9 depressive symptoms included in Diagnostic and Statistical Manual of Mental Disorders (DSM-5) ^19^, interns indicated whether, during the previous 2 weeks, the symptom had bothered them “not at all,” “several days,” “more than half the days,” or “nearly every day.” Each item yields a score of 0 to 3, so that the total score ranges from 0 to 27. A score of 10 or greater on the PHQ-9 was defined as PHQ depression, which has a sensitivity of 93% and a specificity of 88% for the diagnosis of major depressive disorder ^20^. Diagnostic validity of the PHQ-9 is comparable with clinician-administered assessments ^21^.

One to two months prior to internship, subjects completed an online baseline survey, assessing PHQ-9 depressive symptoms, neuroticism (NEO-Five Factor Inventory ^22^), personal history of depression (self-reported yes/no), and early family environment (Risky Families Questionnaire ^23^), along with demographic information. We then contacted the participants via email at months 3, 6, 9 and 12 of their internship year and asked them to complete online quarterly surveys assessing PHQ-9 depressive symptoms and additional information about their internship experience. All surveys were conducted through a secure online website designed to maintain confidentiality, with subjects identified only by numeric IDs. No links between the identification number and the subjects’ identities were maintained. Subjects who completed baseline survey and at least one quarterly survey were included in the analysis, accounting for 86.08% of the total enrolled subjects.

### Genotyping and imputation

We collected DNA from subjects using DNA Genotek Oragene Mailable Tube (OGR-500) ^24^ through the mail. DNA (n = 9,611) was extracted and genotyped on Illumina Infinium CoreExome-24 v1.0 or v1.1 Chip, containing 571,054 and 588,628 SNPs, respectively.

Samples with call rate < 99% (n = 179) or with a sex mismatch between genotype data and reported data (n = 129) were excluded. For 539 duplicated samples, the sample with higher call rate was selected. SNPs were excluded if: on non-autosomal chromosomes, call rate < 0.98 (after sample removal), or MAF < 0.005. Genotype data from v1.0 Chips and v1.1 Chips were then merged and subject duplication was again checked and 11 more duplicates removed. 325,855 SNPs and 8753 samples were considered for further analysis.

We performed linkage disequilibrium (LD)-based pruning (window size 100kb, step size 25 variants, pairwise r^2^ threshold 0.5) which yielded 202,235 SNPs for principal components analysis (PCA) of genotype data using all samples (PLINK version1.9 ^25^). We defined European ancestry samples using the following steps. We plotted the first two PC’s for self-reported European ancestry samples. Based on the plot, to reduce genetic heterogeneity, we included samples that fell within mean PC1±3SD and and mean PC2*±*6SD. We additionally included subjects who did not report ethnicity but whose PC1 and PC2 values were within the range defined above. From this set of European ancestry samples (n = 5,710), we excluded SNPs with Hardy-Weinberg equilibrium P < 10^−6^, leaving 325,249 genotyped SNPs. We defined East Asian ancestry (n = 816) and South Asian ancestry (n = 595) samples with the same procedure, except for East Asian, the sample inclusion range was mean PC1*±*3SD and and mean PC2*±*4.5SD, and for South Asian, it was mean PC1*±*2.5SD and and mean PC2*±*3.5SD (Supplementary Figure 1).

We performed genotype imputation on the Michigan Imputation Server using Minimac3 to phase samples, and the 1000 Genomes Phase 3 data (Version 5, phased by Eagle v2.3) as reference panel.

### Major Depressive Disorder polygenic risk score (MDD-PRS) calculation

To calculate an MDD polygenic risk score of MDD, we used the most recent MDD GWAS summary statistics from a meta-analysis of the Psychiatric Genomics Consortium (PGC) MDD phase 2 and 23andMe containing 135,458 cases and 344,901 controls ^7 26^ (not including Intern Health Study samples). For the MDD-PRS calculation we used 249,588 variants genotyped in our sample (with MAF>=0.1 and outside the MHC region), without applying a PGC/23andMe meta-analysis p-value cutoff. This strategy followed the recommendation from a recent review on PRS calculation ^27^. For comparison, we also calculated MDD-PRS using 115,326 common SNPs (MAF>=0.1) imputed in our sample (imputation quality > 0.9) pruned using LD clumping (500kb window and r^2^<0.1). PRSice V2 ^28^ was used to implement the calculation of MDD-PRS for our intern subjects.

An additive model was used to calculate the polygenic risk score with the formula:

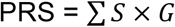

Where S is the PGC2/23andMe GWAS summary statistic effect size for the effect allele, and G = 0,1,2, is the number of effect alleles observed. The MDD-PRS was then mean centered and scaled to a standard deviation of 1.

### Statistical Analysis

Statistical analyses were conducted using R 3.4.4 (The R Foundation, Vienna, AUT) and SAS software Version 9.4 for windows (SAS Institute Inc). We used 5,227 European ancestry samples with completed survey responses and genotype data for the main statistical analyses.

### Association of MDD-PRS with PHQ-9 depressive symptom score

Internship PHQ-9 depressive symptom score was calculated by averaging PHQ-9 score across all available internship assessment. Subjects who reported a PHQ-9 score greater than or equal to 10 in at least one internship assessment were classified as meeting criteria for PHQ depression during internship. We used linear regression to test for association of MDD-PRS with inverse normalized baseline PHQ-9 or internship PHQ-9, adjusting for age, sex and the top 10 principal components of genotype data. We also used logistic regression to assess the association of MDD-PRS with baseline and internship PHQ depression.

To assess whether the effect size of MDD-PRS associating with PHQ-9 or PHQ depression was significantly different at baseline and during internship, we used a linear mixed model or a logistic mixed model to assess the interaction effect between MDD-PRS and internship status, adjusting for main effect of MDD-PRS, internship status, age, sex and the top 10 principal components of genotype data. We included a random intercept term to account for correlation between the repeated measurements within subjects.

### Relationship between MDD-PRS and known risk factors of depression

In a separate analysis, we jointly included the three know risk factors (personal depression history, neuroticism score and early family environment)^12^ as covariates in the models above. Continuous variables including neuroticism score and early family environment were scaled to have a SD of 1.

Mediation analysis was conducted to quantify the proportion of the MDD-PRS and PHQ9 association mediated by known risk factors. R package ‘mediation’ ^29^ was used to fit our data in the following structural equation model:

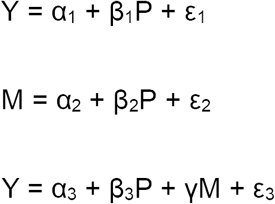

Where Y is the outcome (inverse normalized baseline or internship PHQ-9) and P is the predictor (MDD-PRS). M is the mediator, either one of the known risk factors or the first principal component from a principal component analysis of the three known risk factors. α_i_ (i=1,2,3) and ε_i_ (i=1,2,3) represent the intercepts and errors, respectively. Coefficient β_1_ represents the total effect of the MDD-PRS on PHQ-9, β_2_ represents the effect of MDD-PRS on a known risk factor, β_3_ represents the direct effect of MDD-PRS on PHQ-9 (defined the effect of MDD-PRS that is not accounted for by a known risk factor). γ represents the effect of a known risk factor on PHQ-9. β_2_γ represents the indirect effect of MDD-PRS on PHQ-9 that is mediated by the known risk factor. The total effect of MDD-PRS on PHQ-9 is β_3_ + β_2_γ = β_1_. The mediated proportion β_2_γ / β_1_ is the proportion of the total MDR-PRS effect that is mediated by the known risk factor. To assess the significance of the mediated proportion, we performed 100,000 non-parametric bootstraps to estimate 95% confidence interval and p-value of the mediated proportion test for significant mediation, tested under the null hypothesis of the mediated proportion being zero.

### Differences in PHQ-9 depressive symptoms or PHQ depression proportion by MDD-PRS percentile cutpoint

To test for the ability of higher or lower MDD-PRS to predict different levels of risk of PHQ depression (raw PHQ-9 score >= 10) or PHQ-9, we divided the sample based on an MDD-PRS percentile cut point, including 2.5, 5, 10, 25, 75, 90, 95 and 97.5 percentile cutpoints. At each MDD-PRS percentile cutpoint, we separate the subjects into two subgroups, one is the tail subgroup whose MDD-PRS percentile is lower (for cutpoints<50) or higher (for cutpoints>50) than the cutpoint, the other is the remaining of the subjects. Then we compared the proportion of individuals with PHQ depression in the tail subgroup and the remaining subgroup using a Fisher’s exact test. We also compared the PHQ-9 score in the tail subgroup and the remaining subgroup using a two sample t-test. For pairs of cut points that defined the same size of the sample at the low and high tails of the distribution (for example 5% and 95%), we used a permutation test to assess if the magnitude of the difference between the higher and lower MDD-PRS subgroups differed between the two cutpoints. Specifically, in each of 10,000 permutations, we permuted the MDD-PRS scores for all subjects, reran the Fisher’s exact tests and t-tests for a given pair of cutpoints, and calculated the ratio of the odds ratios ((1/low tail OR) / high tail OR) and the difference of two t-tests statistics (-t-statistic _low tail MDD-PRS-remaining MDD-PRS_) -t-statistic _high tail MDD-PRS-remaining MDD-PRS_). For each observed ratio of the odd ratios, we estimated the two-sided p-value as the number of permutations with a ratio of odds ratios more extreme than either the observed ratio of the odds ratios or 1/(the observed ratio of odds ratios), divided by the number of permutations. For each difference in two t-tests, we calculated the two-sided p-value as the number of permutations with an abs(difference in two t-tests) > abs(observed difference in the two t-tests), divided by the number of permutations.

## URLs

PLINK 1.9: www.cog-genomics.org/plink/1.9/ (Authors : Shaun Purcell, Christopher Chang)

## Data Availability

The de-identified data from this study is available through the Psychiatric Genomics Consortium (PGC): https://www.med.unc.edu/pgc/shared-methods.

### Acknowledgement

We would like to thank the training physicians for taking part in this study. We would also like to thank the research participants and employees of 23andMe, and the participants and researchers of PGC for making this work possible. This project was funded by the National Institute of Mental Health (R01MH101459, PI: Sen).

## Supplementary Materials

**Supplementary Table 1.** MDD Polygenic Risk Score (Generated with Imputed rather than Raw Data) Associations with PHQ-9 Depressive Symptom Scores

| Inclusion of known baseline risk factors* as covariates | Time Point of PHQ-9 score | Beta (SE) |  | p |  |
| --- | --- | --- | --- | --- | --- |
|  |  | MDD-PRS | MDD-PRS x internship stress Interaction** | MDD-PRS | MDD-PRS x internship stress Interaction** |
| Not included | Baseline | 0.042 (0.014) | | $2.96 \times 10^{-3}$ | |
| | Internship | 0.070 (0.012) | | $4.26 \times 10^{-9}$ | |
| | Baseline and internship | 0.041 (0.012) | 0.026 (0.013) | $8.91 \times 10^{-4}$ | 0.03 |
| Included | Baseline | -0.005 (0.013) |  | 0.70 |  |
|  | Internship | 0.025 (0.010) |  | 0.01 |  |
|  | Baseline and internship | -0.003 (0.011) | 0.026 (0.013) | 0.76 | 0.03 |

\* Neuroticism, Personal History of Depression, Early Family Environment
\*\* MDD-PRS x internship stress Interaction term is only available in the model including both baseline and internship data.

**Supplementary Figure 1.**
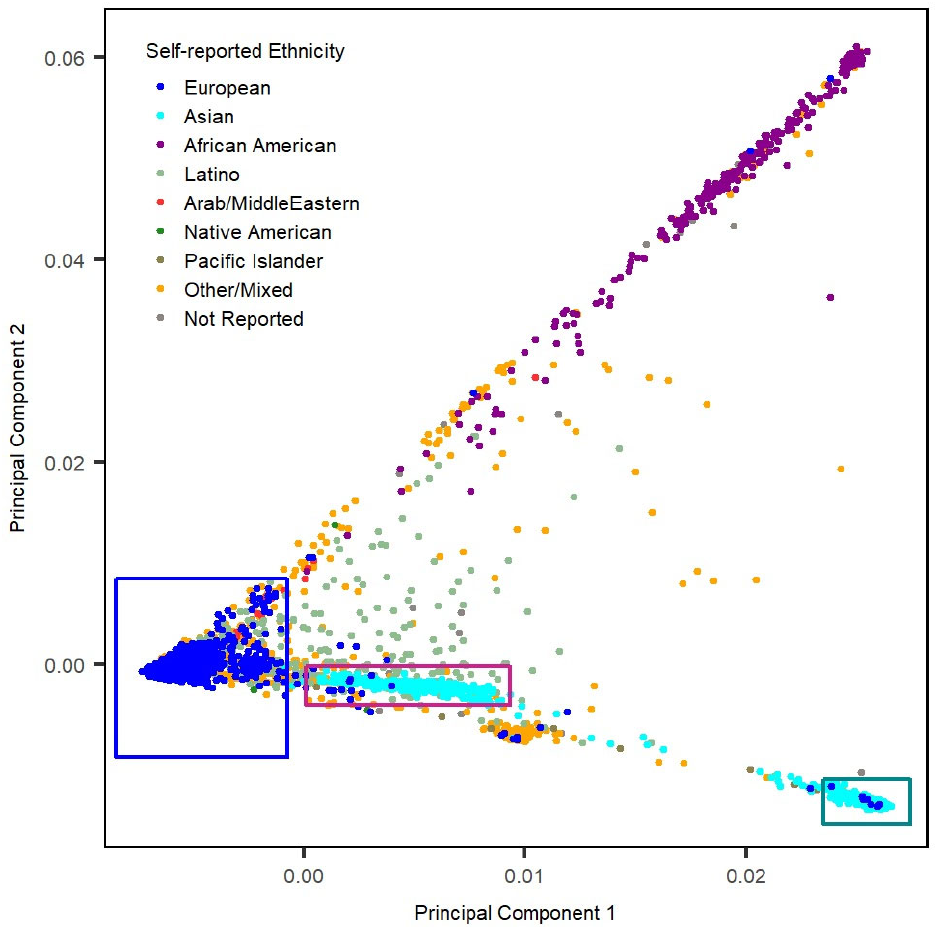
Population Structure Based on the Top Two Principal Component (PC) Analysis of the Intern Health Study. Blue, red and green boxes depicted the analysis inclusion range of European, South Asian and East Asian.

**Supplementary Figure 2.**
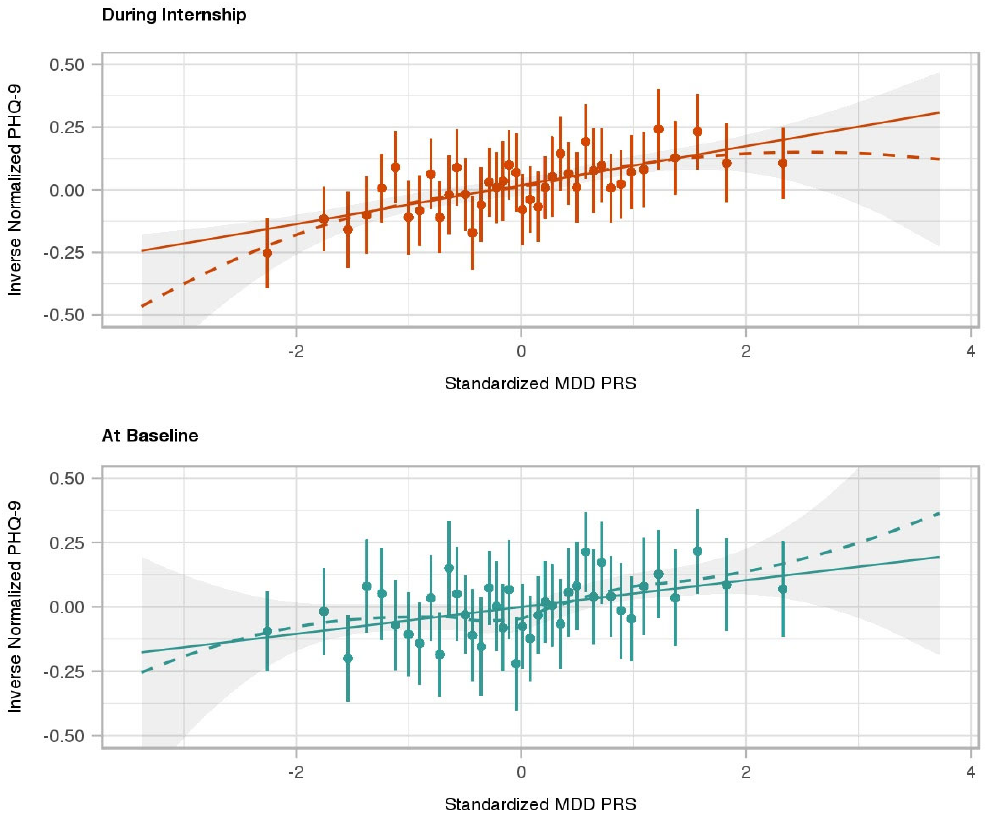
Baseline and Internship PHQ-9 Depressive Symptom Scores by MDD-PRS Group. 5,227 Subjects from Intern Health Study were binned into 40 groups of 2.5% of subjects (n=131 per group) from low to high MDD-PRS (left to right). The 40 x-axis groups are defined by group-wise average standardized MDD-PRS. Average PHQ-9 score of each group at baseline (cyan dots) and during internship (orange dots) are plotted with 95% CI error bar. LOESS fitting line (dash line) shadowed by 95% CI and linear regression fitting line (solid line) were applied to both baseline and internship plots. Optimal span parameter for LOESS regression was selected by generalized cross-validation method.

